# Learning deep representations of enzyme thermal adaptation

**DOI:** 10.1101/2022.03.14.484272

**Authors:** Gang Li, Filip Buric, Jan Zrimec, Sandra Viknander, Jens Nielsen, Aleksej Zelezniak, Martin KM Engqvist

## Abstract

Temperature is a fundamental environmental factor that shapes the evolution of organisms. Learning thermal determinants of protein sequences in evolution thus has profound significance for basic biology, drug discovery, and protein engineering. Here, we use a dataset of over 3 million enzymes labeled with optimal growth temperatures (OGT) of their source organisms to train a deep neural network model (DeepET). The protein-temperature representations learned by DeepET provide a temperature-related statistical summary of protein sequences and capture structural properties that affect thermal stability. For prediction of enzyme optimal catalytic temperatures and protein melting temperatures via a transfer learning approach, our DeepET model outperforms classical regression models trained on rationally designed features and other recent deep-learning-based representations. DeepET thus holds promise for understanding enzyme thermal adaptation and guiding the engineering of thermostable enzymes.

## Introduction

Nature has spent billions of years adapting organisms to various thermal niches, where environmental temperatures range from below -10 °C to over +110 °C ^1^. Since a genome contains all the information required for building and maintaining an organism, the thermal adaptation strategies found in nature are inherently encoded in genomes. In the past decades, much effort has been made to uncover and understand such intrinsic strategies at various levels that include DNA, RNA, proteins and metabolic pathways ^2,3^. Unsurprisingly, most thermal adaptation strategies are clearly reflected at the protein level ^2^, since proteins are involved in almost all cellular functions and are the most temperature sensitive out of all macromolecules ^4–6^. Understanding temperature effects on proteins is also fundamental to basic biology ^5,7,8^, drug discovery ^9^ and protein engineering ^10^. A large portion of studies have thus focused on the temperature effects on protein folding ^5,7,11,12^ and biological functions ^8,13,14^ as well as the combined effects at the systems level ^15–18^. Despite this, it remains unclear how the effects of temperature on a protein are determined by its amino acid sequence.

Although there are many factors that were found to contribute to the thermosensitivity of proteins, including protein length ^12^, amino acid compositions and properties ^19–21^ as well as structural properties ^22–24^, these factors are found to be only weak determinants of the protein thermal properties, such as their unfolding behaviors ^18,25^ and optimal catalytic temperature points ^26^. We hypothesize that by extracting patterns from protein sequences that are related to protein thermal adaptation, we can not only further our understanding of enzyme thermal adaptation, but also provide a rich feature set for many enzyme-related machine learning applications. To this end, in the present study we apply deep-learning to uncover the protein sequence-encoded thermal determinants and learn a predictive representation of enzyme thermal adaptation.

## Results and discussion

### Learning representations of enzyme thermal adaptation

With the assumption that all proteins from an organism should be functional at its optimal growth temperature (OGT), we previously obtained a dataset with 6.5 million enzymes labeled with OGT based on their source organisms ^27^. Here, we removed similar and low-quality sequences and generated a dataset with 3 million enzymes from bacteria, eukaryotes and archaea (Figures 1a,b, Methods). For modeling, we chose the residual neural network architecture ^28^, which has been successfully applied on protein function annotation ^29^. After optimization (Methods, Figures S1-S3), the resulting model contained only 1 residual block with 512 filters (Figure 1c). For model training, one-hot encoded enzyme sequences were used as input and OGT values as output, after which the model could explain ∼60% of the variance in the hold out dataset (Pearson’s *r* = 0.77, *p*-value < 1e-16, Figure 1d). We refer to this model as DeepET hereafter. In DeepET, the network components preceding the Flatten layer can be considered as a feature extractor (Figure 1c), while the last dense layers can be considered as a regressor on top of the above feature extractor. Therefore the values in the Flatten layer (20,480 in total) form a temperature-related representation of input protein sequences (Figure 1c).

**Figure 1.**
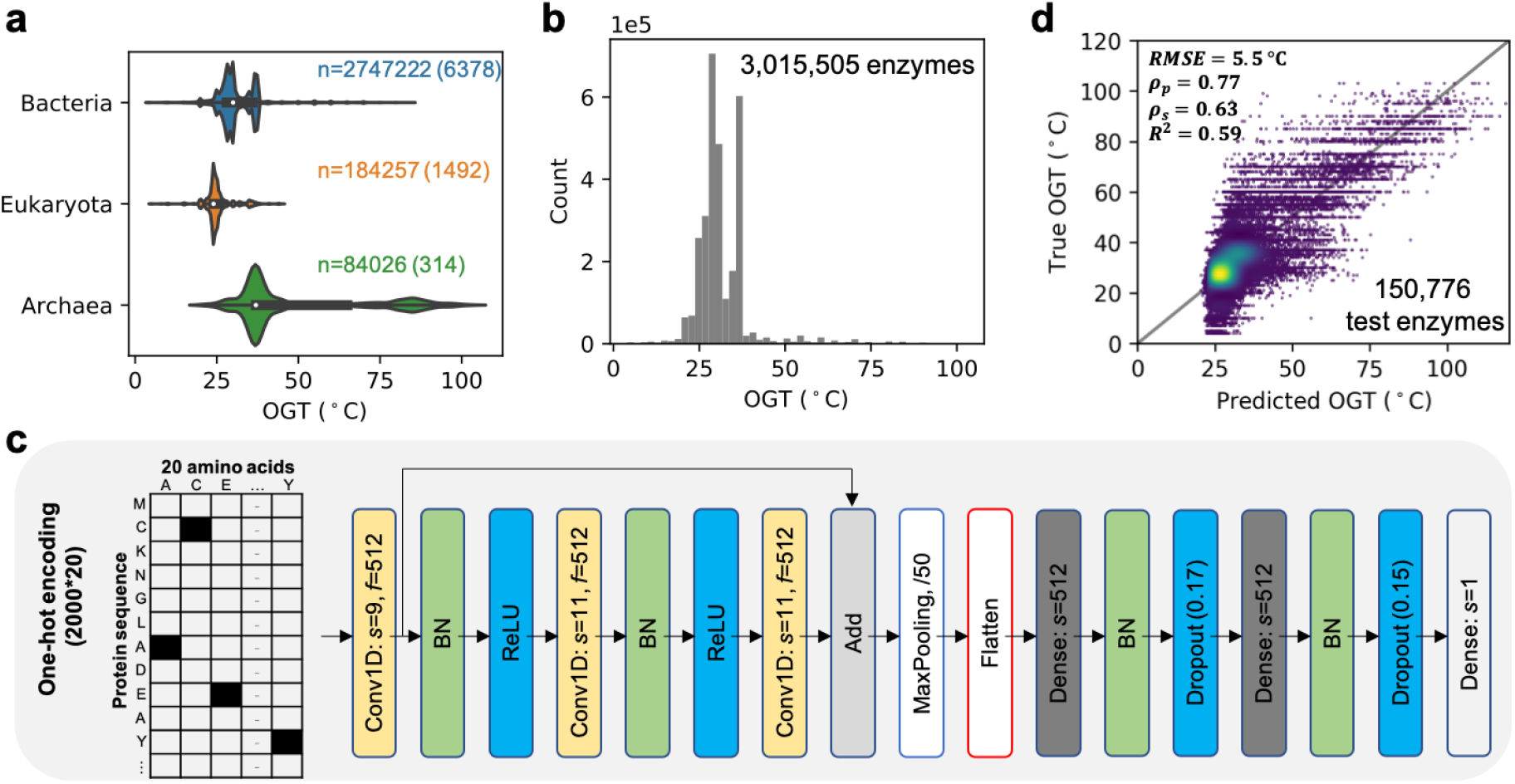
Learning representations of enzyme thermal adaptation with DeepET. (a) OGT distribution of enzymes from three domains, where *n* indicates the number of enzymes from each domain and the number in the parentheses is the number of species where those enzymes are from. (b) The OGT distribution of all enzymes in the training dataset. (c) Optimized architecture of deep neural networks used in this study, where *s* indicates the filter size of convolutional layers and number of nodes for dense layers; *f* indicates the number of filters; /50 in max pooling layer indicates the poolsize and the strides; the floating-point number in a dropout layer indicates the dropout ratio; BN denotes Batch Normalization. (d) Comparison between predicted and true OGT values of enzymes in the hold out dataset. RMSE, root mean squared error; ρ_*p*_, *Pearson*’s correlation coefficient; ρ_*s*_, *Spearman*’s correlation coefficient; *R*^2^, coefficient of determination.

### Transfer learning improves the prediction of protein thermal properties

We next demonstrated the application of DeepET in a transfer learning approach ^30^. In transfer learning, a model pre-trained on a large dataset, such as DeepET, is re-purposed to another similar problem from the same domain with a smaller amount of training samples, by (i) further training and thus fine-tuning certain layers or (ii) resetting their weights and training them from scratch ^31,32^. This is particularly useful for biological datasets since (i) large numbers of biological samples are expensive to collect and (ii) the capacity of classical machine learning models like random forest are usually limited by the availability of relevant features ^26^. Here we chose to predict two critical temperature-related features of proteins: enzyme optimal catalytic temperatures (*T*_opt_), at which the specific activity is maximized, and melting temperatures (*T*_m_), at which there is a 50% possibility that a protein is in a denatured state. For this, two small datasets were collected from literature, one for enzyme *T*_opt_ with 1,902 samples (Figure S4a) ^26^ and another for protein *T*_m_ with 2,506 samples (Figure S4b) ^5^ (Methods). To compare against the deep learning approach, two feature sets were also extracted for classical regression models (Methods): (i) **iFeatures** ^33^, which contains 5,494 protein sequence features, such as amino acid composition and autocorrelation properties; and (ii) **UniRep** ^34^, a multiplicative long-/short-term-memory recurrent neural network (mLSTM) based representation (5,700 features) of protein sequences, which was trained on ∼24 million protein sequences via unsupervised learning. The performance of these two feature sets was tested with six regression models (Figures 2a,b: three best models shown). The deep transfer learning procedure included: (i) training the model shown in Figure 1c **from scratch** (randomly initialized weights); (ii) testing the performance of pre-trained DeepET without any fine-tuning steps (Figures 2a,b: **FrozenAl**l); (iii) freezing convolution layers and fine-tuning the last two dense layers (**FrozenCNN**); (iv) fine-tuning all layers in DeepET (**TuneAll**).

**Figure 2.**
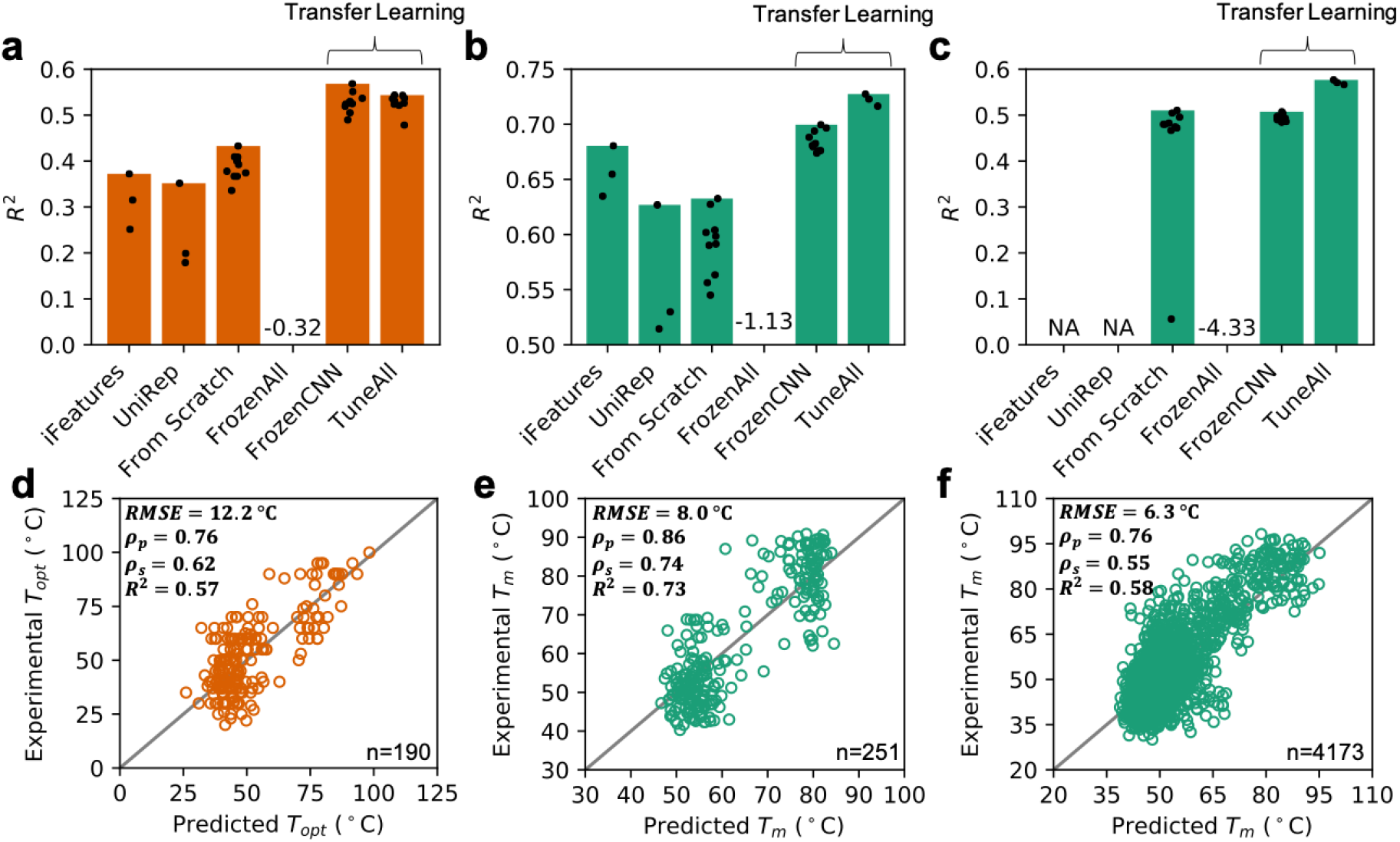
Transfer learning improves the prediction of protein thermal properties. (a-c) *R*^2^ scores of different modeling approaches on hold out datasets which represent 10% of the whole datasets: (a) 190 enzyme optimal catalytic temperatures obtained from ^26^, (b) 251 protein melting temperatures obtained from ^5^ and (c) 4,173 protein melting temperatures obtained from ^7^ (details in Methods section). Bars indicate the maximal *R*^2^ score. iFeatures, the performance of three best classical regression models using features extracted by iFeature ^33^. UniRep, the performance of three best classical regression models using features extracted by UniRep ^34^. From Scratch, the model shown in Figure 1c was trained from scratch (repeated for 10 times). FronzenAll, the pre-trained model was used without any further tuning for prediction. FrozenCNN, froze all layers before Flatten (Figure 1c) and fine-tuned dense layers (repeated 10 times). TuneAll, fine-tuned all layers in the pretrained OGT model (repeated 3 times). (d-f) Comparison between predicted and experimental *T*_opt_/*T*_m_ in hold out datasets. Results of the best model with the highest test *R*^*2*^ score in (a-c) are shown, namely, (d) FrozenCNN, (e) TunaAll and (f) TuneAll. RMSE, root mean squared error; ρ_*p*_, *Pearson*’s correlation coefficient; ρ_*s*_, *Spearman*’s correlation coefficient; *R*^2^, coefficient of determination.

For the tasks of predicting *T*_opt_ and *T*_m_ (Figures 2a,b), DeepET showed superior performance over all other tested strategies when fine-tuning all of its layers (see Methods). For prediction of enzyme *T*_opt_ (Figure 2a), the best model with an *R*^2^ of 0.57 on the hold out dataset was achieved by simply fine-tuning the last two dense layers (Figures 2a,d). This performance is over 50% higher than with the best classical regression models trained on iFeatures or UniRep, and over 30% higher than with the best deep learning model trained from scratch (Figure 2a). The previous best enzyme *T*_opt_ prediction model with an *R*^2^ of 0.61 on the hold out dataset ^26^ was achieved by using amino acid compositions together with OGT as input features. The application of this model is thus limited to native enzymes from microorganisms with known OGT. On the other hand, with DeepET, the new *T*_opt_ model can in principle be applied to any enzyme regardless of organismal sources. For the prediction of *T*_m_ (Figure 2b), melting temperatures of proteins from three microorganisms (*Escherichia coli, Saccharomyces cerevisiae, Thermus thermophilus*) were used^5^. The best model with an *R*^2^ of 0.73 on the hold out dataset was achieved by fine-tuning all layers in DeepET. The performance is 7% higher than with the best model trained on iFeatures, 16% higher than with UniRep and 15% higher than with the model trained from scratch (Figures 2b,e).

The two *T*_opt_ and *T*_m_ datasets (Figures 2a,b) are small and comprise only a few thousand samples, which is why they benefited from the transfer learning approach ^30,35^. Transfer learning may not provide the same benefit for large datasets. Therefore, in the third task, to test if our DeepET network also delivers superior performance for big datasets, we used 41,725 proteins with known melting temperatures from Meltome ^7^ (Figure S4c). Due to the size of this dataset, we could only test the performance of the various deep learning approaches (Figure 2c), since classical models become inefficient to train and optimize ^36^. Surprisingly, fine-tuning all layers still outperforms the model trained from scratch (Figures 2c,f: 13% improvement in *R*^2^).

The results demonstrate that the representations learned by DeepET (values in the Flatten layers, Figure 1c) were predictive in all the above three datasets (Figures 2a-c, FrozenCNN). For the task of predicting enzyme *T*_opt_ (Figure 2a), fine-tuning dense layers achieved similar performance as fine-tuning all layers (Welch’s *t*-test *p*-value = 0.97), meaning that features in the Flatten layer of DeepET are already a collection of informative descriptors for enzyme *T*_opt_. For protein melting temperatures, although the Flatten layer contains informative descriptors for this task, fine-tuning all layers in DeepET showed even better results (Welch’s *t*-test, Figure 2b: *p*-value = 4e-5, Figure 2c, *p*-value = 3e-9).

### Interpreting the sequence determinants of thermostability

Finally, we tested the learned predictive representation of enzyme thermal adaptation by querying the models to identify the specific parts of the protein sequences that were most predictive of optimal catalytic temperature (*T*_*opt*_). For this, we used a perturbation procedure to evaluate the *relevance* of each specific sequence position in relation to the predicted value. Namely, for each protein, we occluded sliding windows of 5 amino acids along its sequence and compared the predictions for all these occlusions with that of the original unoccluded sequence, thus producing a per-residue perturbation or relevance profile for each protein^37,38^ (Methods). The occluded parts of the input protein sequences that yielded a significant deviation in prediction (exceeding ± 2 standard deviations) from the original were regarded as the most relevant for *T*_*opt*_ prediction (Figure 3a). We then checked these relevance profiles against sequence-specific properties that might be salient for prediction: amino acid composition, secondary structure, and protein domains.

**Figure 3.**
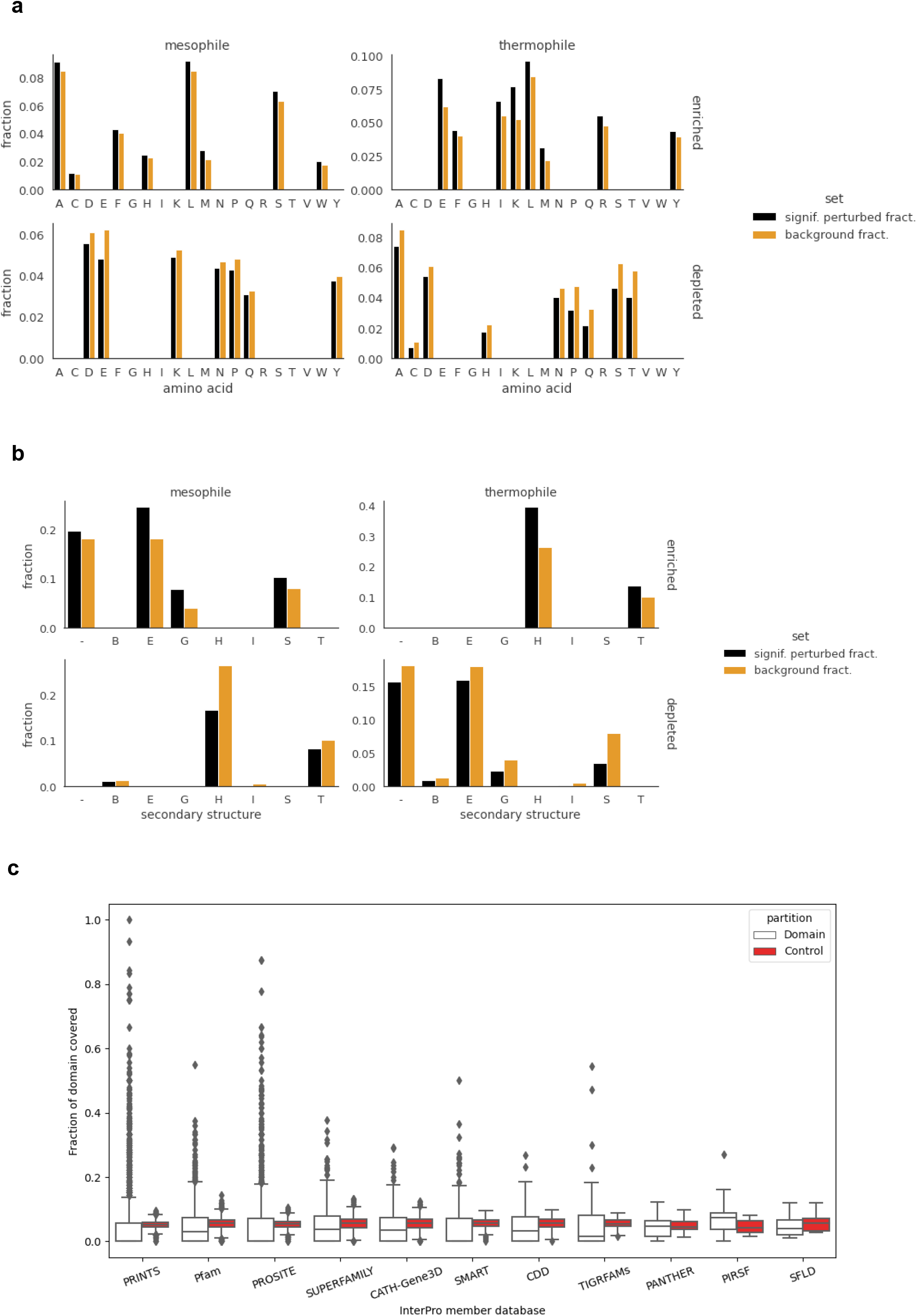
The determinants of thermostability. (a) Enriched and depleted amino acids in the perturbation profiles of mesophiles and thermophiles, showing the most relevant and least relevant amino acids, respectively, towards *T*_*opt*_ prediction. The fractions of amino acids found at significantly (absolute z-score > 2) perturbed positions are compared against the background amino acid distribution. (b) Enriched and depleted secondary structures (in DSSP notation) in the perturbation profiles of mesophiles and thermophiles, showing the most relevant and least relevant structures, respectively, towards *T*_*opt*_ prediction. The fractions of DSSP-annotated secondary structures found at significantly (absolute z-score > 2) perturbed positions are compared against the background amino acid distribution. In DSSP notation, B = isolated β-bridge residues, E = extended strands (parallel or antiparallel β-sheet), G = 3-10 helix, H = ɑ-helix, I = ***π***-helix, S = bend, T= turn, - = coil. (c) Fraction of protein domains covered by significantly perturbed positions, over InterPro domain databases (7,565 domains, 1,227 proteins). PRINTS, PROSITE, and Pfam member databases especially have many domains with high coverage. In total, 219 domains had a coverage of at least 30% across all member databases.

In terms of amino acid composition, the most relevant towards *T*_*opt*_ prediction for mesophiles (OGT 20-45 °C) were Met, Ser, Leu, Ala, Trp, Phe, His, Cys (6 / 8 of which hydrophobic), and conversely, the least relevant were Glu, Pro, Asp, Lys, Asn, Tyr, Gln (6 / 7 of which polar). For thermophiles (OGT > 45 °C), the most relevant were Lys, Glu, Met, Ile, Leu, Arg, Phe, Tyr (5 hydrophobic, 3 polar) and least relevant were Pro, Thr, Ser, Gln, Ala, Cys, His, Asn, Asp (6 / 9 of which polar). See Methods and Figure 3a. These enrichments and depletions are in line with known results^39^ supporting observations that the amounts of (uncharged polar) Cys, Gln, Ser, and Thr are less frequent in thermophiles compared to mesophiles and thus decrease with OGT, while Arg and Tyr increase. Perhaps unintuitively, we find here that Ala relevance decreases between mesophiles and thermophiles, although higher occurrence was noted in thermophiles^40^. There is also some agreement with^21^ in terms of amino acids whose fraction is most correlated with OGT, though we only observe Ile, Tyr, Arg, Glu, and Leu as enriched for our thermophile set. Overall, hydrophobic amino acids generally appear more informative towards prediction, while polar ones are least informative, for both mesophilic and thermophilic groups, which corroborates protein hydrophobicity as an indicator of thermal adaptation^39^. As a commonality between mesophiles and thermophiles, we found that Met, Leu, and Phe occurrence (in decreasing order of enrichment for both) to be determinant for predictions.

In terms of secondary structure (as predicted per-residue by DSSP from PDB files), the most relevant towards prediction for mesophiles were strands (E), 3-10 helices (G), bends (S), and coils (-) (Figure 3b). Least relevant were ɑ-helices (H), ***π***-helices (I), turns (T), and isolated β-bridge residues (B). For thermophiles, most relevant were ɑ-helices (H) and turns (T). Thus, while for mesophiles all major secondary structure types are observed to factor towards prediction, only helices and turns are the most determinant for thermophiles. This is in line with the known increase in helical content with higher temperature adaptations^41^, due to its importance in stabilization. The increase in relevance of helices also fits with the enrichment of Arg and depletion of Cys, His, and Pro relevance for thermophile prediction observed here, as these are known to be favored and disfavored for helix formation, respectively^41^.

To assess whether certain protein domains are more salient for prediction, we measured the overlap of the relevance profiles with domains from the InterPro database^42^, as the fraction of domain positions covered by significant (absolute z-score > 2) perturbation values. Control consisted of measuring the coverage outside the domain, by dividing the outside sequence into windows with the same length as the domain, then taking the average coverage across these (Methods). To ensure proper control, the search was limited to domains no longer than half of the protein sequence, a total of 7,565 domains across 1,227 proteins. While the median relevance profile coverage of domains did not greatly differ from control, the distributions of domain coverage fractions were quite wide and heavy-tailed across the InterPro member databases, with many domains clearly higher than control (Figure 3c). In total, 219 domains had a coverage of at least 30% (a cutoff chosen to be distinctly larger than any control). To get an overview of these domains, we took the GO terms associated with the protein domains (retrieved from InterPro) and produced GO slims using the Generic GO Subset (Methods). While the GO slims for mesophilic enzymes spanned a diverse range of biological processes, including metabolic processes, stress response, protein transport, immune involvement, and cell adhesion, thermophile terms were limited to metabolic processes and response to stress (Tables S3 and S4). Thus, with increased temperature adaptation, the domains responsible for these latter functions are more determinative for the prediction of *T*_*opt*_.

## Conclusion

Here we presented DeepET, a deep learning model that learns temperature-related representations of protein sequences. We demonstrated that these representations are highly useful for the prediction of enzyme catalytic temperature optima and protein melting temperatures, by using a transfer learning approach. Our base model was trained to predict optimal growth temperature with a high *R*^*2*^ from 3 million enzyme sequences across all three domains of life. The model was then re-purposed via fine-tuning to predict optimal enzyme catalytic temperature, as well as melting temperature, both of which showed good performance. As the base DeepET model was trained from sequence alone, the transfer approach is more applicable than previous deep learning approaches for optimal catalytic temperature prediction ^26^, which rely on extracted sequence features (amino acid composition) and optimal growth temperature as input, the latter of which may not be available.

The good performance on the transfer learning prediction tasks suggests that the representations indeed capture and provide a statistical summary of the enzyme thermal adaptation strategies from nature. To get insights into the sequence factors that are informative for the prediction of optimal catalytic temperature, we performed a perturbation analysis by exhaustively occluding sequences with a sliding window to measure the impact on the predicted value, then analyzing the properties of the most relevant sequence positions thus perturbed. We found a large overlap with known determinants of thermostability in terms of amino acid composition and secondary structure, both generally and when distinguishing between mesophiles and thermophiles. This gives greater confidence in the quality and general applicability of the learned DeepET features through transfer learning or data mining for sequence properties and patterns. Checking the relevant positions against protein domains, we saw that while the associated biological processes of domains present in mesophiles covered a wider range, domains thus found in thermophiles were limited to metabolic processes and response to stress, hinting at the adaptations of these enzymes for higher temperatures.

Given these recapitulations of known primary and secondary protein structure determinants of thermostability by DeepET’s features, which were learned by the model from sequence alone, and the observed shift in model-relevant domains between mesophiles and thermophiles, the use of the DeepET is a promising avenue towards elucidating the physical mechanisms that convey enzymes resistance to extreme temperatures. Future work will therefore focus on further interpreting DeepET and its learned representations both using *in silico* analyses, and in a biological context, to deepen our understanding of enzyme thermal adaptation.

## Methods

### Dataset with OGT labeled enzyme sequences

6,270,107 enzyme sequences with unique Uniprot IDs were collected from the previous study ^27^. After removal of sequences that were (1) longer than 2,000; or (2) shorter than 100; or (3) with any non-standard amino acids, there were 6,141,006 enzyme sequences left. Then the cd-hit algorithm (-c 0.95, -T 20, -M 0 and other parameters as default) ^43^ was applied to cluster those sequences into 3,016,273 clusters. Only the representative sequence of each cluster was used for the next step, to keep the resulting sequences diverse. At last, 768 sequences were removed since they were present in the *T*_opt_ dataset (see next section) by matching Uniprot IDs. In the end, a dataset with 3,015,505 enzyme sequences from microorganisms with known optimal growth temperatures was obtained. The dataset was randomly split into training (2,864,729 enzymes) and test (150,776 enzymes) datasets based on a 95-5 ratio.

### Dataset with enzyme optimal catalytic temperatures (*T*_opt_)

This dataset was taken from ^26^, which contains 1,902 enzymes with known *T*_opt_ collected from BRENDA ^44^. The dataset was randomly split into training (1,712 enzymes) and test (190 enzymes) datasets based on a 90-10 ratio.

### First Dataset with protein melting temperatures (*T*_m_)

Leuenberger *et al*. ^5^ experimentally measured melting temperatures for more than 8,000 proteins from four species *(Escherichia coli, Saccharomyces cerevisiae, Thermus thermophilus*, and human cells). In this study, 2,506 proteins from three microorganisms (*E. coli, S. cerevisiae, T. thermophilus*) with experimentally measured *T*_m_ were obtained, after removal of ones with sequences that were (1) longer than 2,000; or (2) shorter than 100; or (3) with any non-standard amino acids. The dataset was randomly split into training (2,255 enzymes) and test (251 enzymes) datasets based on a 90-10 ratio.

### Second Dataset with protein melting temperatures (*T*_m_)

Jarzab *et al*. ^7^ reported melting temperatures for 48,000 proteins from 13 species, ranging from archaea to humans. We first collected all *T*_m_ values from all 77 data sets and corresponding sequence IDs. Only proteins with an existing uniprot ID and protein sequence in the Uniprot database were considered. After removal of sequences that were (1) longer than 2,000; or (2) shorter than 100; or (3) with any non-standard amino acids, a dataset with 41,725 proteins was obtained. For those proteins with multiple *T*_m_ values, the mean value was used. The dataset was randomly split into training (37,552 enzymes) and test (4,173 enzymes) datasets based on a 90-10 ratio.

### Deep neural networks

In the present study, we used Residual networks ^28^, with the model architecture (Figure S1) similar to those that had been applied to protein functional annotation previously ^29^. It contains 1-3 residual block(s) followed by 2 fully connected (FC) layers (Figure S1b). Batch Normalization ^45^ was applied after all layers; Weight dropout ^46^ was applied after FC layers and max-pooling ^47^ was applied after the last residual blocks. The Adam optimizer ^48^ with mean squared error loss function and ReLU activation function ^49^ with uniform ^50^ weight initialization were used.

### Hyper-parameter optimization

Two small OGT datasets with 10,000 samples each were used for tuning hyper-parameters: (1) the first one was randomly sampled from the OGT training dataset (Figure S2a); (2) the second one was sampled in a way that the resulting samples showed a uniform OGT distribution (Figure S2b). Each dataset was randomly split into training (90%) and validation (10%) datasets. The hyper-parameters were tuned using values randomly sampled from the defined parameter spaces (Table S1). Around 100-200 parameter sets were randomly sampled and tested. The best hyper-parameter set was chosen based on the one with the lowest validation loss on each small OGT dataset. Then the model with these two hyper-parameter sets was tested with the big OGT training dataset (2,864,729 enzymes). This dataset was further split into training (95%) and validation (5%) datasets. After manually tuning a few hyper-parameters, the best model with the lowest validation loss was chosen as the final hyper-parameter set (Figure S3).

### Feature extraction for enzymes in *T*_opt_ and two protein *T*_m_ datasets

A set of 5,494 rationally designed features was extracted with iFeature ^33^. These features included k-mer compositions (AAC, 20 features; DPC, 400), composition of k-spaced amino acid pairs (CKSAAP, 2400), dipeptide deviation from expected mean (DDE, 400), grouped amino acid composition (GAAC, 5), composition of k-spaced amino acid group pairs (CKSAAGP, 150), grouped dipeptide composition (GDPC, 25), grouped tripeptide composition (GTPC, 125), Moran autocorrelation (Moran, 240), Geary autocorrelation (Geary, 240), normalized Moreau-Broto (NMBroto, 240), composition-transition-distribution (CTDC, 39; CTDT, 39; CTDD, 195), conjoint triad (CTriad, 343), conjoint k-spaced triad (KSCTriad, 343), pseudo-amino acid composition (PAAC, 50), amphiphilic PAAC (APAAC, 80), sequence-order-coupling number (SOCNumber, 60) and quasi-sequence-order descriptors (QSOrder, 100).

### UniRep

A representation with 1900×3 features was extracted for each protein sequence with the previously published deep learning model UniRep ^34^, which is a Multiplicative Long-Short-Term-Memory (mLSTM) Recurrent Neural Network that was trained on the UniRef50 dataset ^51^.

### Supervised classical ML methods

Two linear regression algorithms BayesianRidge and Elastic Net as well as three non-nonlinear algorithms Decision Tree, Random Forest and Support Vector Machine were evaluated on each feature set (iFeatures and UniRep). Input features were firstly scaled to a standard normal distribution by 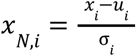, where *x*_*i*_ is the values of feature i of all samples, *u*_*i*_ and *σ*_*i*_ are the mean and standard deviation of *x*_*i*_, respectively. This was done by taking all samples, including train and test datasets together. The training dataset was further randomly split into training and validation datasets. The validation dataset was used to tune the hyper-parameters via a greedy search approach. The optimized model was tested on the held out test dataset and the *R*^2^ score was calculated. In Figure 3a and 3b, the *R*^2^ score of three best regression models were shown for iFeature and UniRep. All machine learning analyses in this section were performed with scikit-learn (v0.20.3) ^52^ using default settings.

### Relevance profile analysis

The sequence perturbation study was performed on a set of 1554 enzymes, a subset of the *T*_*opt*_ dataset. Relevance profiles were obtained for each sequence by sliding a 5-amino-acid-long occlusion window on the sequence, 1 amino acid at a time (thus resulting in overlapping windows). For each window position, a *T*_*opt*_ prediction was obtained and the perturbation or relevance score was calculated as (*prediction*_*occluded*_ - *prediction*_*wt*_) / *prediction*_*wt*_. As the sliding window position was bounded by the sequence, in order to obtain a perturbation vector of the same length as the sequence (and thus, a relevance score for each amino acid), the sequence was flanked by 2 repeats of the terminal amino acids. A moving average was then performed on the resulting relevance score vector for each protein. The width of the occlusion window was chosen to be small and match short secondary structure features. The impact of the occlusion width was tested by performing the perturbation procedure with windows of length 2, 5, 10, and 20, which respectively represent 0.42%, 1%, 2%, and 4.2% of the average sequence length in the set, 477. The resulting profiles showed large overlaps (FIgure S5) and the choice of width had no impact on the resulting set of significantly covered protein domains (Figure S6).

The amino acid enrichment of relevance profiles was assessed by performing one-sided hypergeometric tests between the background amino acid counts in all sequences and the counts of amino acids occurring at significant (absolute z-score > 2) relevance profile positions, to test for both overrepresentation (enrichment) and underrepresentation (depletion) of amino acids. A p-value threshold of 0.05 was set for significance. Cryophiles (OGT < 20 °C, 5 sequences) were excluded due to very low counts. The remaining set included 1220 mesophiles and 323 thermophiles. An analogous procedure was performed to assess relevance of secondary structure, starting from per-residue sequence annotations obtained with DSSP 3 from PDB files. Due to either lack of PDB entries or structural errors within the files, the structural annotation set only included 874 mesophiles and 279 thermophiles. As a sanity check, positions where no annotation was available appeared as underrepresented (depleted) and removed from results.

The InterPro database (retrieved 2021-06-24) was filtered to only domains at most half of the length of the protein, to ensure balance when performing control. The per-domain control consisted in taking the sequence *S*_*out*_ outside of a given domain and counting the number of significantly perturbed (absolute z-score > 2) positions. This number was divided by the number of windows in *S*_*out*_ of the same length as the domain, to give an expected count corresponding to repeatedly randomly sampling subsequences the same length as the domain. The final control coverage fraction was taken as the above average divided by the domain length.

GO slims were produced starting from the GO terms provided for each domain in the InterPro database and the Generic GO Subset provided by the GO Consortium (version 2021-08-21) ^53,54^. We selected domains that had at least 30% of their length covered by significantly perturbed (absolute z-score > 2) positions. The processing was performed using the Python packages GOATOOLS 1.1.6 ^55^and obonet 0.3.0. The full list of terms of GO slims is given in Tables S3 and S4.

### Software

Python v3.6 (www.python.org) scripts were used for the computations and data analysis, using the packages NumPy 1.18.1 ^56^, SciPy 1.6.2 ^57^, tensorflow 1.14 ^58^, keras 2.2.4, Biopython 1.76 ^59^, PySpark 3.1.2. The code and data are available at Zenodo (https://doi.org/10.5281/zenodo.6351465).

## Author contributions

GL and MKME conceptualized the project; GL, FB, JZ, SV, AZ and MKME designed the research pipeline; GL, FB, JZ and SV performed computations; GL, JZ, SV, MKME and AZ interpreted results; AZ and JN provided funding acquisition; GL and FB wrote the draft manuscript; All authors carried out revisions on the final manuscript version.

## Acknowledgements

GL and JN have received funding from the European Union’s Horizon 2020 research and innovation program under the Marie Skłodowska-Curie program, project PAcMEN (grant agreement No 722287). JN also acknowledges funding from the Novo Nordisk Foundation (grant no. NNF10CC1016517), the Knut and Alice Wallenberg Foundation. The study was supported by SciLifeLab funding and the Swedish Research council (Vetenskapsrådet) starting grant no. 2019-05356. AZ was supported by the Marius Jakulis Jason foundation and JZ by the Slovenian Research Agency (ARRS) grant no. J2-3060 and Public Scholarship, Development, Disability and Maintenance Fund of the Republic of Slovenia grant no. 11013-9/2021-2. The computations were performed on resources at Chalmers Center for Computational Science and Engineering (C3SE) provided by the Swedish National Infrastructure for Computing (SNIC).

