## Supplementary information for "Learning deep representations of enzyme thermal adaptation"

#### Table of contents

Supplementary figures … p.2 - p.7

Supplementary tables … p.8 - p.11

Supplementary references … p.12

### Supplementary Figures

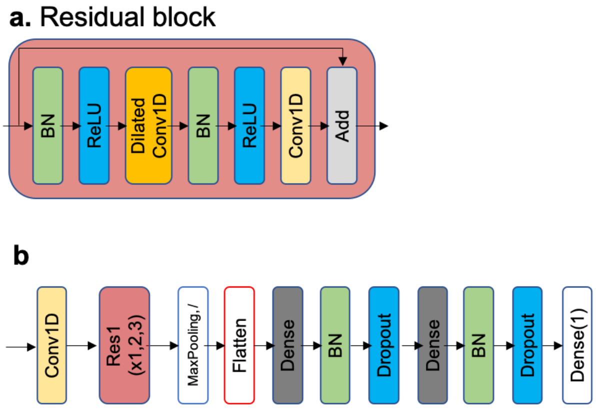


**Figure S1.** The ResNet based model architecture used in this study. (a) the residual block, taken from (*1*) (b) the model architecture. It takes one-hot encoded protein sequences as input. The number of residual blocks (Res1) was treated as a hyperparameter varying from 1-3.

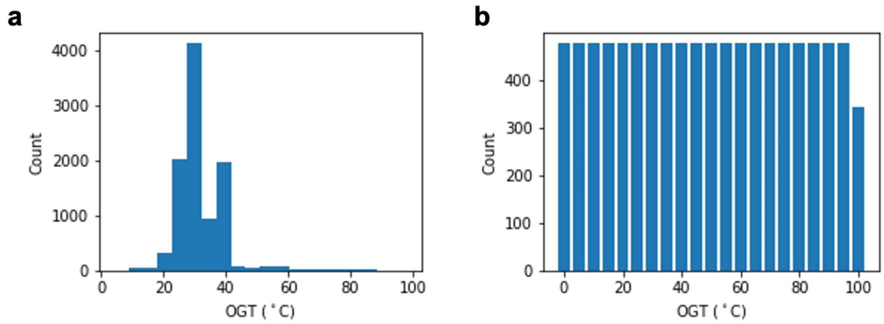


**Figure S2.** OGT datasets for hyper-parameter optimization. (a) the distribution of OGT values of enzymes randomly sampled from the original training dataset. (b) a uniformly distributed dataset was sampled from the original training dataset. There are 10,000 enzymes in each dataset.

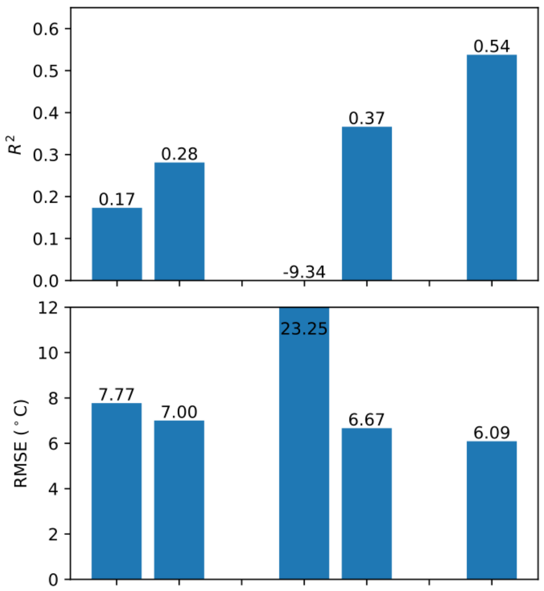


| | | | | | |
|---|---|---|---|---|---|
| **Distribution** | ori | uni | ori | uni | uni |
| **filters** | 128 | 512 | 512 | 512 | 512 |
| **lr** | 5e-4 | 1e-4 | 5e-4 | 1e-4 | 1e-4 |
| **Batch size** | 32 | 32 | 32 | 32 | 128 |
| **Dense2** | 512 | 64 | 512 | 512 | 512 |

**Figure S3.** Validation metrics from hyper-parameter tuning results. The upper and lower panel are the R2 score and root mean squared error on the validation dataset. The description under the figure is variables tuned. **Distribution**, if the hyper-parameter set is achieved on a dataset with original distribution (ori) or uniform distribution as shown in Figure S2; **filters**, the number of filters used in all convolution layers; **lr**, learning rate; **Dense2**, the number of nodes in the second fully-connected layer (Figure S1). The hyper-parameter sets of the first two bars were obtained by a random search approach on two datasets as shown in Figure S2. Then the number of filters and the size of the Dense2 were increased to 512, respectively (3rd and 4th bar). The bad scores shown in the 3rd bar were due to the training not being finished in 7-days. Since the model architecture optimised on the uniformly distributed dataset shows better performance on big OGT validation dataset, it was further tuned by increasing the **batch size** from 32 to 128. In the end, the hyper-parameters used for the last bar were considered as the final hyper-parameters (Detailed list is in Table S2).

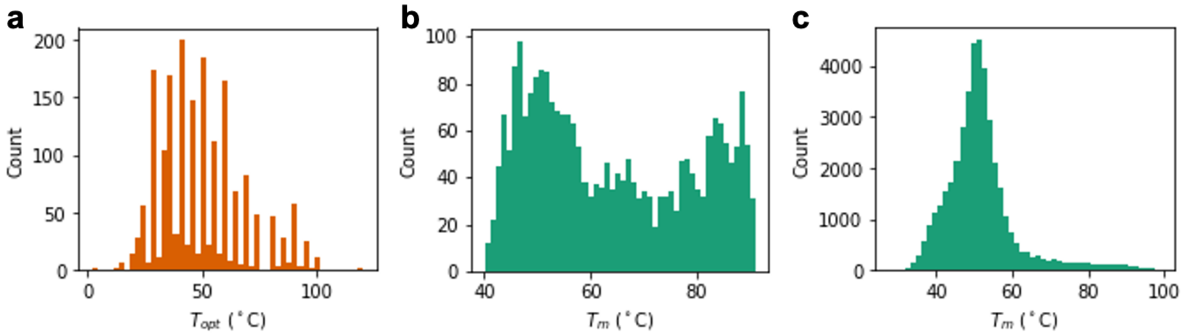


**Figure S4.** Distributions of (a) enzyme $T_{opt}$ from (*2*), protein melting temperatures from Leuenberger P et al (*3*) and Jarzab A et al (*4*). See details in the Methods section.

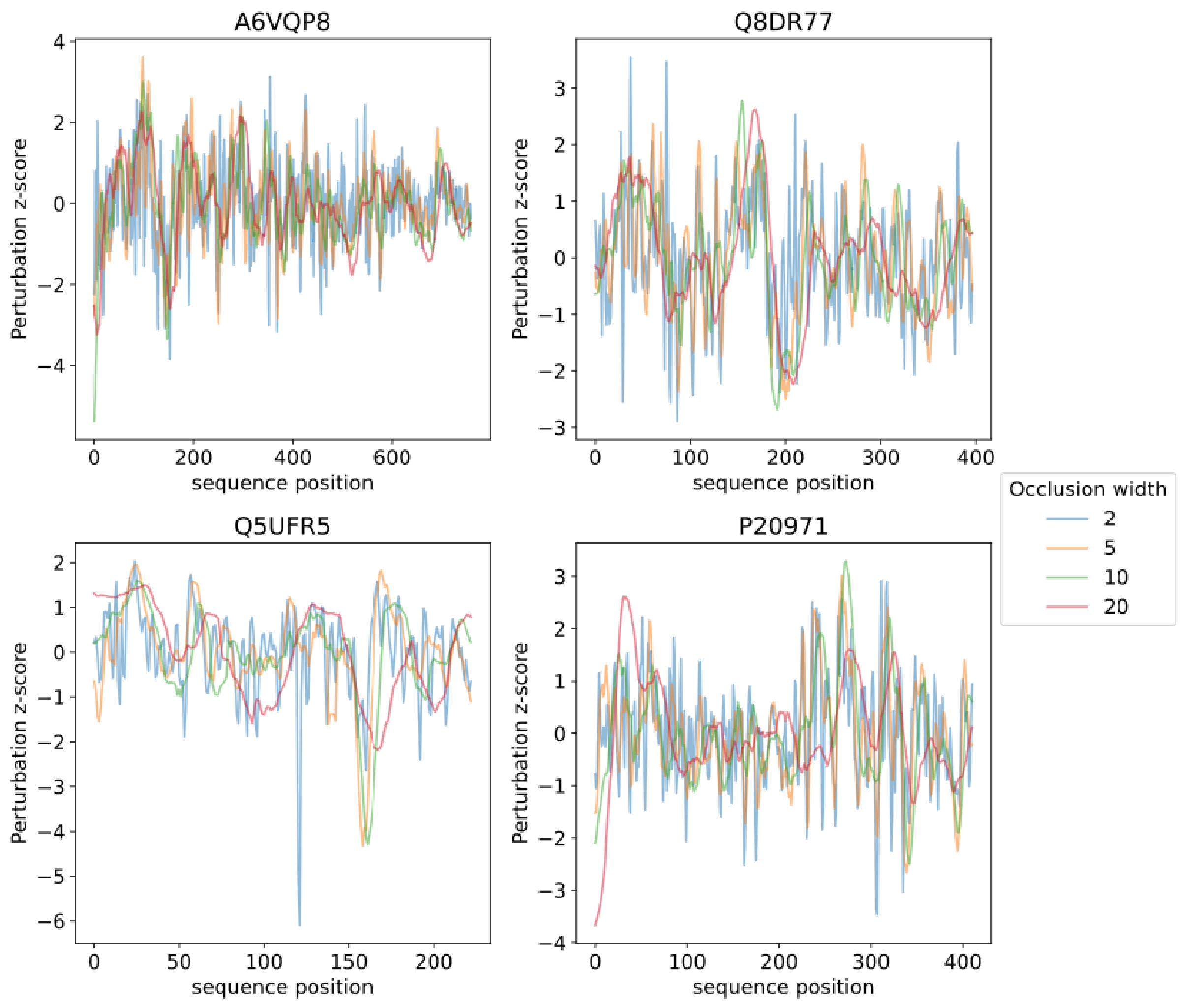


**Figure S5.** Comparison of perturbation profiles using different occlusion widths, for 4 randomly selected sequences (UniProt IDs given as subfigure titles), showing the overall large similarity when using different widths.

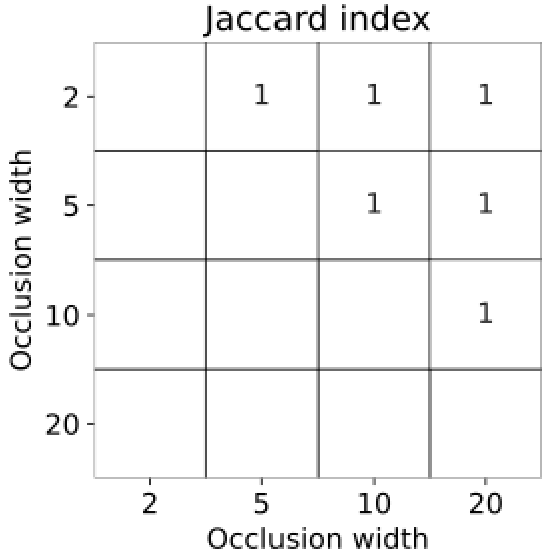


**Figure S6.** The choice of occlusion window width has no impact on the set of protein domains covered by significant perturbation profile positions. Indeed, the sets resulting from different occlusion window widths overlap perfectly, as shown here by the Jaccard index of 1 between these (showing only the upper triangle, as this measure is symmetric).

### Supplementary Tables

**Table S1.** Hyper parameter space used for random search search

| Name | Space | Applied to layers |
|---|---|---|
| filters | [32, 64, 128, 256, 512] | All convolution layers have the same number of filters |
| kernel size | [3, 7, 9, 11, 21, 31] | All convolution layers, they have different kernel sizes |
| dilation (ResBlock) | [1, 2, 3, 5] | The first conv1d layer in ResBlock |
| pool size | [2, 4, 8, 20, 30, 40] | The max pooling layer, pool size is equal to strides |
| Dense 1 | [256, 512, 1024] | First dense layer |
| Dense 2 | [128, 256, 512] | Second dense layer |
| Dropout | (0,0.5) | Two dense layers, they have different dropout values |
| *lr* | [1e-4, 5e-4, 1e-3] | |
| mbatch | [32, 64, 128, 256] | |
| Number of Residual blocks | [1, 2, 3] | |

**Table S2.** Optimised hyper-parameters

| | | OriDist (Figure S2a) | UniDist(Figure S2b) |
|---|---|---|---|
| | filters | 128 | 512 |
| | Kernel size 1 | 7 | 9 |
| Residual Block 1 | Kernel size 21 | 7 | 21 |
| | Kernel size 22 | 7 | 11 |
| | dilation2 | 1 | 1 |
| Residual Block 2 | Kernel size 31 | 31 | NA |
| | Kernel size 32 | 21 | NA |
| | dilation3 | 3 | NA |
| | Pool size (=strides) | 30 | 50 |
| | dense1 | 512 | 512 |
| | Drop out 1 | 0.35 | 0.17 |
| | dense2 | 512 | 64 → **512** |
| | Drop out 2 | 0.37 | 0.15 |
| | lr | 5e-4 | 1e-4 |
| | mbatch | 32 | 32 → **128** |
| | Total weights | 5.8M | 19.0M → 19.2M |

**Table S3**. Biological process GO slims for the most relevant domains for $T_{opt}$ prediction of mesophilic and thermophilic enzymes.

| GO ID | Mesophiles | Thermophiles |
|---|---|---|
| GO:0009058 | biosynthetic process | |
| GO:0044281 | small molecule metabolic process | small molecule metabolic process |
| GO:0006259 | DNA metabolic process | DNA metabolic process |
| GO:0002376 | immune system process | |
| GO:0007155 | cell adhesion | |
| GO:0034641 | cellular nitrogen compound metabolic process | |
| GO:0006605 | protein targeting | |
| GO:0009056 | catabolic process | catabolic process |
| GO:0015031 | protein transport | |
| GO:0005975 | carbohydrate metabolic process | carbohydrate metabolic process |
| GO:0006091 | generation of precursor metabolites and energy | |
| GO:0006950 | response to stress | response to stress |
| GO:0006629 | lipid metabolic process | |
| GO:0006520 | cellular amino acid metabolic process | cellular amino acid metabolic process |
| GO:0006399 | | tRNA metabolic process |

**Table S4**. Molecular function GO slims for the most relevant domains for $T_{opt}$ prediction of mesophilic and thermophilic enzymes.

| GO ID | Mesophiles | Thermophiles |
|---|---|---|
| GO:0016491 | oxidoreductase activity | oxidoreductase activity |
| GO:0016779 | nucleotidyltransferase activity | |
| GO:0016791 | phosphatase activity | phosphatase activity |
| GO:0016301 | kinase activity | kinase activity |
| GO:0008233 | peptidase activity | peptidase activity |
| GO:0016829 | lyase activity | |
| GO:0016765 | transferase activity, transferring alkyl or aryl (other than methyl) groups | |
| GO:0043167 | ion binding | ion binding |
| GO:0003677 | DNA binding | |
| GO:0016798 | hydrolase activity, acting on glycosyl bonds | hydrolase activity, acting on glycosyl bonds |
| GO:0016810 | hydrolase activity, acting on carbon-nitrogen (but not peptide) bonds | |
| GO:0008168 | | methyltransferase activity |
| GO:0016874 | | ligase activity |
| GO:0016853 | | isomerase activity |
